# Identifying conservation priorities of a pantropical plant lineage: a case study in Scleria (Cyperaceae)

**DOI:** 10.1101/2024.06.21.600016

**Authors:** Javier Galán Díaz, Steven P. Bachman, Félix Forest, Marcial Escudero, Hannah Rotton, Isabel Larridon

**Author notes:** Declaration Of Authorship: JGD and IL originally formulated the idea. JGD, HR and IL gathered and curated the data. JGD, SPB, FF and ME performed statistical analyses. JGD, SPB, FF, ME, HR and IL wrote the manuscript. JGD and IL acquired funding.

## Abstract

*Scleria* is a pantropical genus of annual and perennial herbs and the sixth largest genus in the Cyperaceae family with around 261 species. In this study, we produced preliminary extinction risk assessments for the ∼30% of *Scleria* species that do not yet have a global Red List assessment and followed the Evolutionarily Distinct and Globally Endangered (EDGE2) and Ecologically Distinct and Globally Endangered (EoDGE) protocols to identify evolutionary and ecologically unique *Scleria* species at greatest risk of extinction and hotspots of rare and endangered species. Our results indicate that 38 of the 78 *Scleria* species not yet included in the Red List, and 26% of species in the genus, are potentially threatened with extinction. The risk of extinction is not equally distributed across the phylogeny, and the Afrotropics and the Neotropics accumulate most threatened species. Eleven ecoregions mostly from four African (Madagascar, D.R. Congo, Zambia and Tanzania) and two South American (Brazil, Venezuela) countries accumulate almost half of *Scleria* species and stand out in terms of their sum of EDGE2 scores. Phylogenetic and functional distinctiveness metrics were largely uncorrelated, and the EcoDGE metric mostly points towards South American countries as reservoirs of ecologically distinctive and endangered species: Brazil, Venezuela, Bolivia, Peru, Colombia, Guyana and Dominican Republic. Recent methodological advances in the identification of species at-risk of extinction and the novel EDGE2 framework emerge as powerful tools to identify conservation priorities.

## Introduction

*Scleria* P.J.Bergius (1765: 142), commonly known as nut rushes or razor grasses, is the sixth largest genus in the Cyperaceae family with around 260 species (Bauters et al. 2016; Larridon et al. 2021a), and the only genus of tribe Sclerieae Wight & Arn. (subfamily Cyperoideae) (Larridon 2022). *Scleria* has four subgenera and 17 sections (Bauters et al. 2016, 2018, 2019). Most species occur in tropical areas but some occupy more temperate climatic regions (Larridon et al. 2021b). *Scleria* is a functionally diverse genus that includes annual and perennial herbs, the latter often using stoloniferous rhizomes or tubers (Bauters et al. 2016; Galán Díaz et al. 2019). Recent studies have disentangled the evolutionary relationships among *Scleria* species (Bauters et al. 2016, 2018) and identified major dispersal and niche shift events (Larridon et al. 2021b). Still, we lack a global assessment of species and world regions of remarkable interest for *Scleria* conservation.

In the current context of rapid global change, there is a need to adopt new tools to assess biodiversity and ecosystems to set conservation priorities (Cowell et al. 2022). Over the last two decades, the use of phylogenetic and functional indices has emerged as a powerful approach to support conservation assessments by identifying at-risk species and regions of great evolutionary and ecological uniqueness (Kondratyeva et al. 2019). The Evolutionarily Distinct and Globally Endangered (EDGE) metric can be used to rank species conservation priorities where species’ evolutionary distinctiveness (ED) is used as a surrogate of their irreplaceability, and species’ probability of extinction (GE) is used as a proxy of their vulnerability (Isaac et al. 2007). It has been successfully implemented in different groups such as corals (Huang 2012), mammals and amphibians (Safi et al. 2013), gymnosperms (Forest et al. 2018) and tetrapods (Gumbs et al. 2018). The EDGE metric has been recently revised to include new advances (EDGE2; Gumbs et al. 2023): (i) species’ evolutionary distinctiveness (ED2) now considers the extinction risk of their close relatives; (ii) it allows the incorporation of uncertainty in the phylogeny and extinction risk. The latter allows the use of phylogenies which have been expanded using non-molecular information (e.g., see Ramos-Gutiérrez et al. 2023), and provides a flexible framework to assess Not Evaluated and preliminary assessed species rather than only depending on species published in the Red List of Threatened Species of the International Union for Conservation of Nature (IUCN) (e.g., Walker et al. 2023).

Machine learning approaches constitute efficient tools to produce extinction risk assessments and their implementation in different plant groups has provided promising results (Zizka et al. 2022; Bachman et al. 2023; Walker et al. 2023). The latest version of the Red List includes ∼18% (62,666 species) of all known vascular plant species (IUCN 2023). In the case of *Scleria*, there are extinction risk assessments available for 183 species (26 of which are not yet publicly available in the latest version of the Red List), which accounts for 70% of the genus. Recent taxonomic revisions of the genus (Bauters et al. 2016, 2018, 2019; Galán Díaz et al. 2019) and the availability of occurrence datasets curated using expert knowledge (Larridon et al. 2021b) can help overcome some of the limitations and challenges that arise from using herbarium collections to support preliminary extinction risk assessments (Nic Lughadha et al. 2019). The genus *Scleria* is therefore a good study system to train machine learning algorithms and produce preliminary extinction risk assessments.

Finally, the phylogenetic and functional components of biodiversity are not necessarily correlated, especially when considering functional attributes which are highly dependent on the environment (Losos 2008). *Scleria* is an ecologically diverse clade that includes species with variable habits, from tiny annuals with small nutlets and fibrous roots to climbers or stout perennials with big buoyant propagules (Galán Díaz et al. 2019). The variation in functional traits across the genus *Scleria* therefore warrants independent evaluation. In this regard, the EDGE2 framework can incorporate species functional distinctiveness (FUD) as a measure of ecological irreplaceability. This metric is termed Ecologically Distinct and Globally Endangered (EcoDGE) (Hidasi-Neto et al. 2015).

Here, (i) we first produced preliminary extinction risk assessments for the ∼30% of *Scleria* species that do not yet have a global Red List assessment, and then (ii) followed the EDGE2 protocol to identify evolutionary and ecologically distinct *Scleria* species at greatest risk of extinction and conservation priority areas. For this, we used extinction risk assessments from the global Red List and the most comprehensive phylogeny of *Scleria* to date (Larridon et al. 2021b), as well as newly compiled occurrence and traits datasets.

## Material and Methods

### Taxonomy and species occurrences

We used the World Checklist of Vascular Plants (WCVP; Govaerts et al. 2021) and the latest taxonomic revisions of the genus *Scleria* P.J.Bergius (Bauters et al. 2016, 2019) to compile a dataset of 261 accepted species. Data relating to infraspecific taxa were incorporated at the species level. Species names and authorities are available in supplementary material Appendix 1.

Occurrence data was compiled using observations from the Global Biodiversity Information Facility (GBIF 2023), Red List (IUCN 2023), research-grade identifications from iNaturalist (accessed 13/08/2023) and records for collections from BR, K, GENT, L, MO, NY, P, US and WAG which were georeferenced using Google Earth. For each species, we filtered observations following several steps: (1) we removed duplicates and retained one observation per 1km^2^ raster cell, (2) filtered observations based on the native ranges at the ‘botanical country’ scale, according to the WCVP, and (3) excluded observations 1.5 times outside the interquartile climatic range using the mean annual temperature and annual precipitation BIOCLIM variables (Booth et al. 2014). The final occurrence database included 22,759 observations from 248 species. We projected locality data using the Equal Earth map projection (šavrič et al. 2019).

### Traits measurements and calculation

We measured maximum height, maximum blade length, maximum blade width, nutlet length and nutlet width from 1,254 specimens housed at Royal Botanic Gardens, Kew (K) and the Muséum National d’Histoire Naturelle in Paris (P). For species missing in these herbaria, we completed the dataset using information from protologues and descriptions from regional floras (e.g., Davidse et al. 1994, Simpson and Koyama 1998, Le Roux 2015). This information was used to estimate three traits informative of plant ecological strategies (Westoby 1998): maximum height (i.e., distance from the top inflorescence to the ground), leaf area and size of the propagule (i.e., nutlet volume). We estimated leaf area and nutlet volume using the formula of an ellipse and an ellipsoid, respectively. We also considered plant longevity (annual/perennial).

### Extinction risk data and preliminary assessments

We downloaded 157 extinction risk assessments from the Red List (version 2023-1) and considered the extinction risk category of other 26 species that have assessments in preparation. To supplement the 183 species with global (published and in preparation) Red List assessments we carried out preliminary assessments and predictions for the 67 Data Deficient and Not Evaluated species for which occurrence data was available so that we had an extinction risk assessment for all species. The extinct species *S. chevalieri* J.Raynal was excluded from the analyses. We followed two approaches:

i. We estimated species extinction risk categories under IUCN Red List criterion B using the function ‘ConBatch’ from the R package ‘rCAT’ (Moat 2017). Given a dataset of occurrences, this function provides preliminary extinction risk categories based on species’ extent of occurrence (EOO). We did not consider the species’ area of occupancy (AOO) because it can lead to an overestimation of extinction risk when using occurrence data derived from herbarium records (Nic Lughadha et al. 2019). To assess the accuracy of this method, we ran ‘ConBatch’ on the species included in the Red List and compared the actual assessments with the predicted categories.
ii. We implemented a machine learning algorithm (random forest, henceforth RF) using as predictors: EOO, AOO, mean of latitudinal range, elevation, minimum human population density in 2020 (HPD; CIESIN 2018), human footprint index in 2013 (HFI; Venter et al. 2018), proportion of observations located in protected areas (UNEP-WCMC and IUCN 2023), mean annual temperature, minimum temperature of the coldest month, temperature annual range, annual precipitation, precipitation of the driest month and precipitation seasonality. All predictors were resampled at 30 seconds. For HPD and HFI, we calculated the average index value within a 5 km circular buffer of each unique occurrence point. We used 10 repeats of 5-fold cross-validation to train and evaluate the model and retained 20% of data for external validation.

### EDGE2 and EcoDGE calculation and species lists

We computed two metrics that combine species extinction risk with their evolutionary or functional distinctiveness to prioritize at-risk *Scleria* species and world regions of special interest for conservation: the Evolutionarily Distinct and Globally Endangered metric (EDGE2) (Gumbs et al. 2023) and the Ecologically Distinct and Globally Endangered metric (EcoDGE) (Hidasi-Neto et al. 2015). To compute ED2, we used the latest phylogenetic inference of the genus from Larridon et al. (2021a), which included 136 species. We used the R package ‘randtip’ (Ramos-Gutiérrez et al. 2023) to impute 111 species missing in the phylogeny but for which infrageneric information (i.e., section) was available. We excluded 14 species for which infrageneric information was not available. Trees were expanded following several criteria: species were imputed in their sections at random, the probability of branch selection was set as equiprobable, and the stem branch was considered as a candidate for binding. This step was repeated to generate a distribution of 500 randomly imputed phylogenetic trees.

To calculate FUD, we adapted the EcoDGE framework proposed by Hidasi-Neto et al. (2015) to the new EDGE2 protocol following Griffith et al. (2022). We generated a dendrogram by calculating pairwise species dissimilarities with a generalized Gower’s distance matrix (Gower 1971) using the four traits (i.e., longevity, maximum height, leaf area and size of the propagule) and an unweighted pair group method with arithmetic mean (UPGMA). We repeated the dendrogram construction 11 times (all possible combinations of two, three and four traits) to reduce the impact of specific traits on the results. Finally, we multiplied FUD scores by 100 to balance the weighting of the two component values of the EcoDGE metric.

The EDGE2 and EcoDGE protocols allow the incorporation of uncertainty in the quantification of extinction probabilities by associating each Red List category to a distribution of extinction probabilities based on a 50-year time horizon (Mooers et al. 2008). For species not included in the Red List and classified as threatened by the preliminary assessments, the extinction risk component (GE) was randomly drawn from CR, VU and EN categories. For species classified as non-threatened, GE values were randomly withdrawn from those of NT and LC categories.

Finally, for each species, we calculated a distribution of EDGE2 and EcoDGE scores (N=500) which were used to sort *Scleria* species in three lists as proposed by Gumbs et al. (2023): a ‘priority or main EDGE2 species list’ that includes threatened species whose distribution of EDGE2 scores rank above the median of the entire genus at least 95% of the times; a ‘borderline EDGE2 species list’ that includes threatened species whose distribution of EDGE2 scores rank above the median of the entire genus at least 80% of the times; and a ‘watch EDGE2 species list’ that includes nonthreatened species whose distribution of EDGE2 scores rank above the median of the entire genus at least 95% of the times. We further discuss the results in the context of the countries and world terrestrial ecoregions (Olson et al. 2001), which are good descriptors of plant distribution (Smith et al. 2018) and can be used as informative units for conservation planning (Dinerstein et al. 2017).

All analyses were performed in R (version 4.3.2).

## Results

### Preliminary assessments

rCAT and RF yielded different results (Table 1; supplementary material Appendix 2). Under criterion B, rCAT classified 53 out of the 67 non-assessed species as threatened according to their EOO (44 CR, 5 EN, 4 VU, 4 NT, 10 LC). *Scleria* rCAT assessments correctly classified 79.31% of threatened species (high sensitivity) and 91.39% of non-threatened species (high specificity). RF classified 39 species as threatened and 28 as non-threatened. It achieved an accuracy of 97.22%, correctly classifying 96.55% of non-threatened species and all threatened species (higher sensitivity and specificity than rCAT). The single most important predictor for RF was EOO, followed by AOO. Because of its greater performance, we retained the results of RF for the EDGE2 and EcoDGE calculations. Combined Red List results (published, unpublished and preliminary) are available in supplementary material Appendix 1.

**Table 1.** Confusion matrices of preliminary assessments. To evaluate the performance of rCAT (Moat 2017), we compared the IUCN category of the species included in the Red List with the output of the function ‘ConBatch’. rCAT preliminary assessments follow IUCN Red List criterion B estimated from species’ EOO. The accuracy of the random forest model was tested on a subset of 20% of observations left for external validation.

| Method |  | predicted values |  | observed values |
| --- | --- | --- | --- | --- |
|  |  | non-threatened | threatened |  |
| rCAT EOO | non-threatened | 138 | 13 |  |
|  | threatened | 6 | 29 |  |
| Random forest | non-threatened | 28 | 1 |  |
|  | threatened | 0 | 7 |  |

Considering the published (Red List version 2023-1) and preliminary (RF) results, we found that the Afrotropics and the Neotropics had the highest number and proportion of threatened species (Fig. 1a). The sections with the greatest proportion of threatened species were *Hypoporum* (41.54%), followed by Abortivae, Browniae, Schizolepsis and Foveolidia (28.57%) (Fig. 1b).

**Fig. 1.**
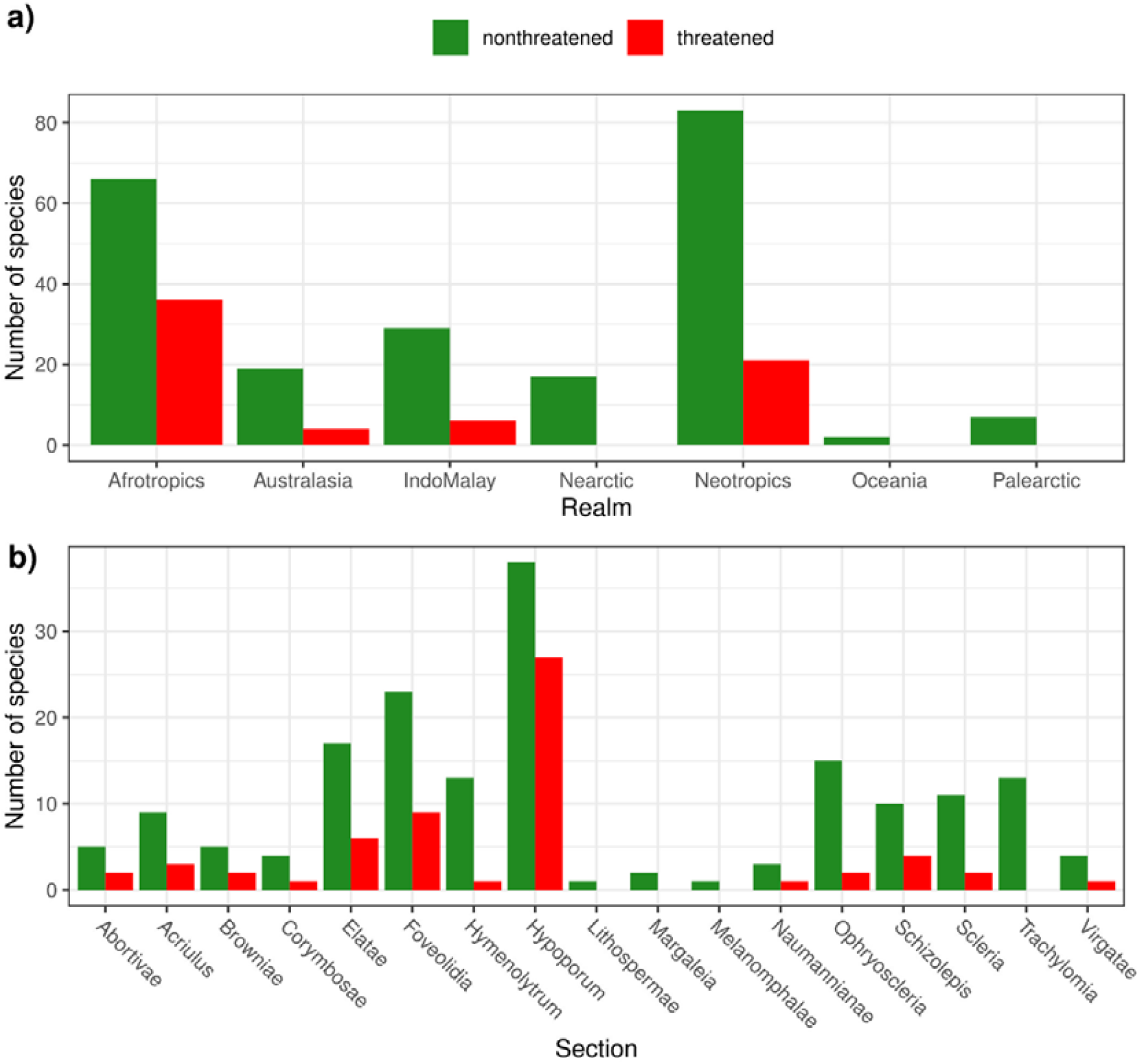
Number of threatened and non-threatened *Scleria* species grouped by **(a)** Realm (Olson et al. 2001) and **(b)** Section. Threatened refers to species classified as CR, EN and VU in the Red List, as well as DD and NE species classified as threatened by the random forest approach. Non-threatened refers to species classified as LC and NT in the Red List, as well as DD and NE species classified as non-threatened by the random forest approach.

### Evolutionarily and ecologically distinct species at greatest risk of extinction

The phylogenetic diversity (PD) across trees was 900.96 ± 24.01 MY (mean ± SD). Species with the highest mean ED2 scores were *Scleria corymbosa, Scleria lithosperma* and *Scleria melanomphala* with 22.28, 20.89 and 19.04 MY, respectively. The species with the greatest mean EDGE2 score was *Scleria porphyrocarpa* with 3.97 MY of avertable expected PD loss, followed by *Scleria zambesica* (2.53 MY), *Scleria pulchella* (2.25 MY) and *Scleria madagascariensis* (2.04 MY).

We detected 45 EDGE2 *Scleria* species (17.24% of species in the genus). Safeguarding these species, we would secure 68.24% (41.49 MY) of avertable expected PD loss in a 50-year time horizon. The main list included 23 species (Table 2), of which 17 belong to section *Hypoporum*, three to section *Acriulus*, two to section *Abortivae* and one to *Corymbosae*. The EDGE2 borderline and watch lists included 17 and 5 species respectively (supplementary material Appendix 2). The watch list included widespread and locally abundant species that belong to evolutionary distinctive sections such as *S. lithosperma, S. melanomphala* and *S. tonkinensis*.

**Table 2.** EDGE2 main list, i.e., threatened species that scored above the median EDGE2 score of the entire genus at least 95% of the time. Species are ranked based on their EDGE2 score. IUCN RL: Red List category of published species and preliminary assessment of Not Evaluated species (i.e., Threatened). EDGE2: Evolutionarily Distinct and Globally Endangered metric, ED2: evolutionary distinctiveness.

| Scientific name | Section | IUCN RL | EDGE2<br>(mean ± sd) | ED2<br>(mean ± sd) |
| --- | --- | --- | --- | --- |
| *S. porphyrocarpa* E.A.Rob. | *Corymbosae* | Threatened | 3.97 ± 6.26 | 8.89 ± 8.27 |
| *S. zambesica* E.A.Rob. | *Hypoporum* | Threatened | 2.53 ± 1.46 | 5.1 ± 0.87 |
| *S. pulchella* Ridl. | *Hypoporum* | CR | 2.25 ± 0.45 | 2.44 ± 0.42 |
| *S. madagascariensis* Boeckeler | *Abortivae* | EN | 2.04 ± 0.66 | 4.62 ± 1.01 |
| *S. ankaratrensis* Bauters | *Hypoporum* | DD (Threatened) | 1.45 ± 0.82 | 2.77 ± 0.48 |
| *S. maypurensis* Bauters | *Hypoporum* | CR | 1.45 ± 0.35 | 1.65 ± 0.34 |
| *S. pachyrrhyncha* Nelmes | *Acriulus* | EN | 1.37 ± 1.68 | 2.81 ± 3.32 |
| *S. induta* Turrill | *Acriulus* | Threatened | 1.35 ± 2.43 | 2.73 ± 3.53 |
| *S. thwaitesiana* Boeckeler | *Acriulus* | Threatened | 1.2 ± 2.4 | 2.56 ± 3.33 |
| *S. fulvipilosa* E.A.Rob. | *Hypoporum* | EN | 1.07 ± 0.3 | 2.31 ± 0.38 |
| *S. richardsiae* E.A.Rob. | *Hypoporum* | EN | 0.96 ± 0.27 | 2.09 ± 0.37 |
| *S. nusbaumeri* Bauters | *Abortivae* | Threatened | 0.96 ± 1.34 | 1.92 ± 1.76 |
| *S. pantadenia* Meganck & Bauters | *Hypoporum* | VU | 0.94 ± 0.26 | 4.17 ± 0.75 |
| *S. tricristata* Meganck & Bauters | *Hypoporum* | EN | 0.84 ± 0.29 | 1.85 ± 0.44 |
| *S. calcicola* E.A.Rob. | *Hypoporum* | EN | 0.82 ± 0.22 | 1.74 ± 0.29 |
| *S. burchellii* C.B.Clarke | *Hypoporum* | EN | 0.81 ± 0.26 | 1.83 ± 0.39 |
| *S. liberica* Bauters | *Hypoporum* | VU | 0.59 ± 0.16 | 2.57 ± 0.43 |
| *S. polyrhiza* E.A.Rob. | *Hypoporum* | EN | 0.57 ± 0.16 | 1.24 ± 0.21 |
| *S. mongomoensis* Bauters | *Hypoporum* | EN | 0.41 ± 0.16 | 0.87 ± 0.27 |
| *S. pedicellata* Bauters | *Hypoporum* | EN | 0.4 ± 0.16 | 0.88 ± 0.27 |
| *S. pseudohispidioides* Bauters | *Hypoporum* | VU | 0.31 ± 0.1 | 1.39 ± 0.29 |
| *S. tricholepis* Nelmes | *Hypoporum* | VU | 0.28 ± 0.08 | 1.18 ± 0.24 |
| *S. spicata* (Spreng.) J.F.Macbr. | *Hypoporum* | VU | 0.27 ± 0.08 | 1.14 ± 0.23 |

*Scleria* species with the highest mean FUD scores were *Scleria skutchii*, the tallest species in the genus, and *Scleria depressa*, the species with the largest nutlet. Overall, 6 of 10 species with the highest FUD scores belonged to the sections *Ophryoscleria* and *Schizolepis* which include stout perennials with large leaf areas and climbers. The species with the greatest mean EcoDGE score was *Scleria porphyrocarpa*, followed by *Scleria tropicalis, Scleria williamsii* and *Scleria chlorantha*. We found 38 EcoDGE species, 6 in the main list (Table 3) and 32 in the borderline EDGE2 species list (supplementary material Appendix 3). These species (14.56% of species of the genus) account for 64.28% of avertable expected functional diversity loss in the genus in a 50-year time horizon.

**Table 3.**
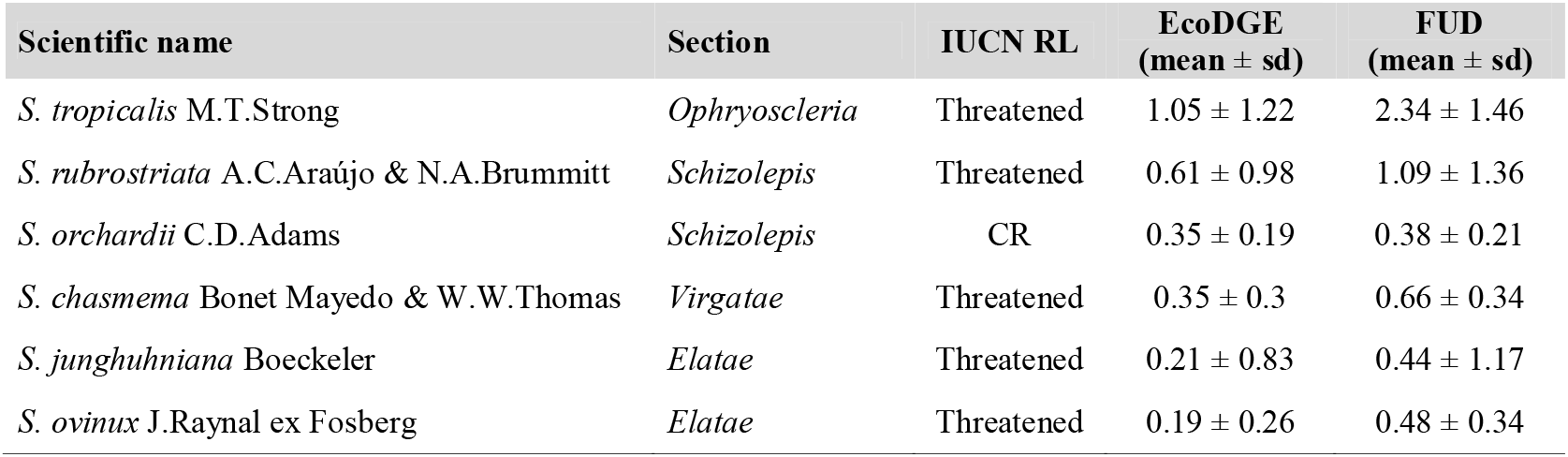
EcoDGE main list, i.e., threatened species that scored above the median EcoDGE score of the entire genus at least 95% of the time. Species are ranked based on their EcoDGE score. IUCN RL: Red List category of published species and preliminary assessment of Not Evaluated species (i.e., Threatened). EcoDGE: Ecologically Distinct and Globally Endangered metric, FUD: functional distinctiveness.

| Scientific name | Section | IUCN RL | EcoDGE<br>(mean ± sd) | FUD<br>(mean ± sd) |
| --- | --- | --- | --- | --- |
| *S. tropicalis* M.T.Strong | *Ophryoscleria* | Threatened | 1.05 ± 1.22 | 2.34 ± 1.46 |
| *S. rubrostriata* A.C.Araújo & N.A.Brummitt | *Schizolepis* | Threatened | 0.61 ± 0.98 | 1.09 ± 1.36 |
| *S. orchardii* C.D.Adams | *Schizolepis* | CR | 0.35 ± 0.19 | 0.38 ± 0.21 |
| *S. chasmema* Bonet Mayedo & W.W.Thomas | *Virgatae* | Threatened | 0.35 ± 0.3 | 0.66 ± 0.34 |
| *S. junghuhniana* Boeckeler | *Elatae* | Threatened | 0.21 ± 0.83 | 0.44 ± 1.17 |
| *S. ovinux* J.Raynal ex Fosberg | *Elatae* | Threatened | 0.19 ± 0.26 | 0.48 ± 0.34 |

The evolutionary distinctiveness (ED2) and functional distinctiveness (FUD) metrics of *Scleria* were not correlated (F_1,237_=0.03, p=0.86; Fig. 2).

**Fig. 2.**
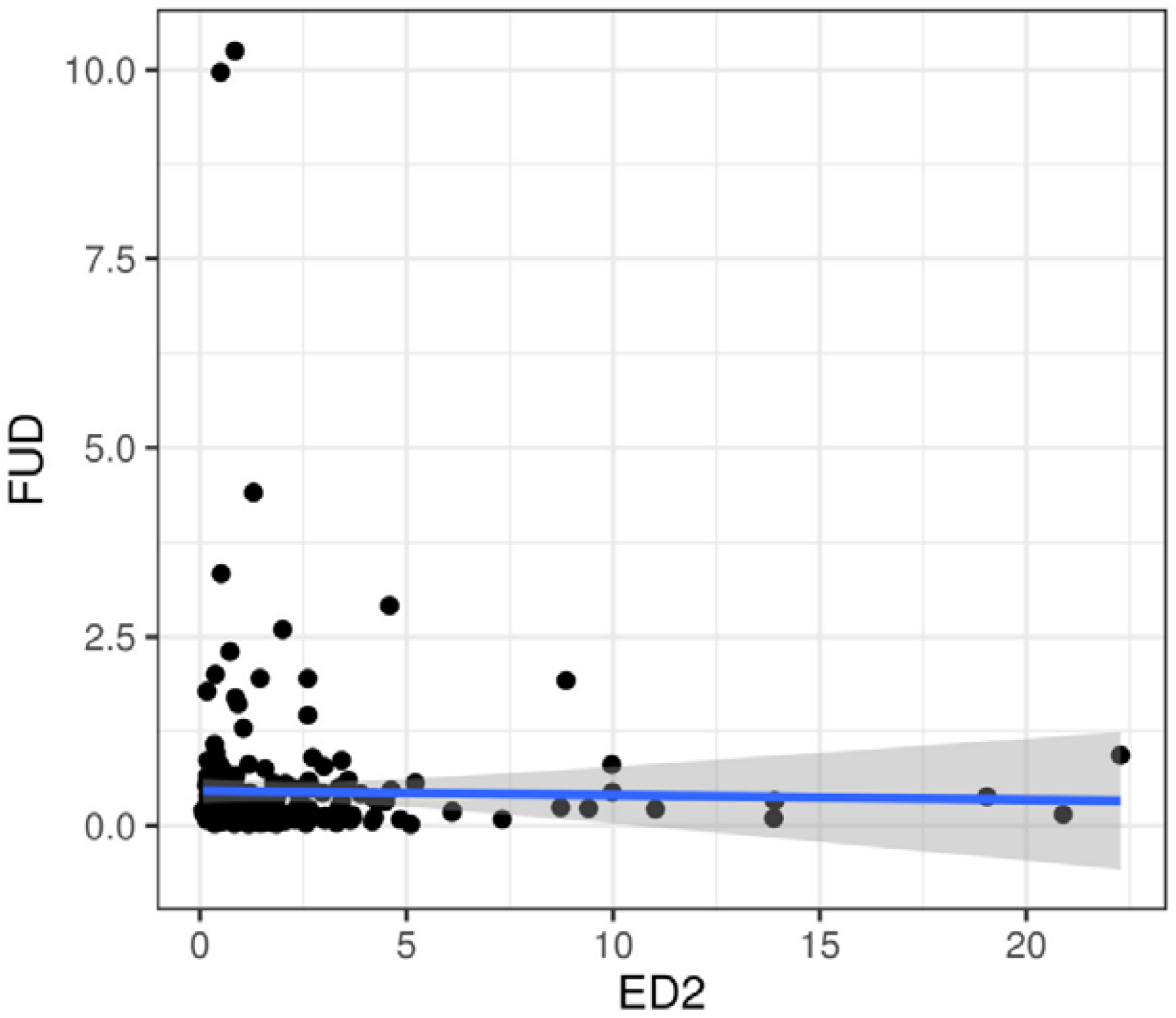
Relationship between the two irreplaceability surrogates of *Scleria* used in this study: evolutionary distinctiveness (ED2) and functional distinctiveness (FUD). F(1,237)=0.03, p=0.86.

### Regions of special interest for conservation

The countries with the highest sum of EDGE2 scores and the greatest number of listed EDGE2 species were Madagascar, Democratic Republic of the Congo, Brazil, Zambia and Tanzania (Table 4). The ecoregions with the highest sum of EDGE2 scores and the greatest number of listed EDGE2 species were the Central Zambezian Miombo woodlands, Madagascar subhumid forests, Cerrado, East Sudanian savanna, Llanos, Guinean forest-savanna mosaic and Madagascar lowland forests (Table 5; Fig. 3b).

**Table 4.**
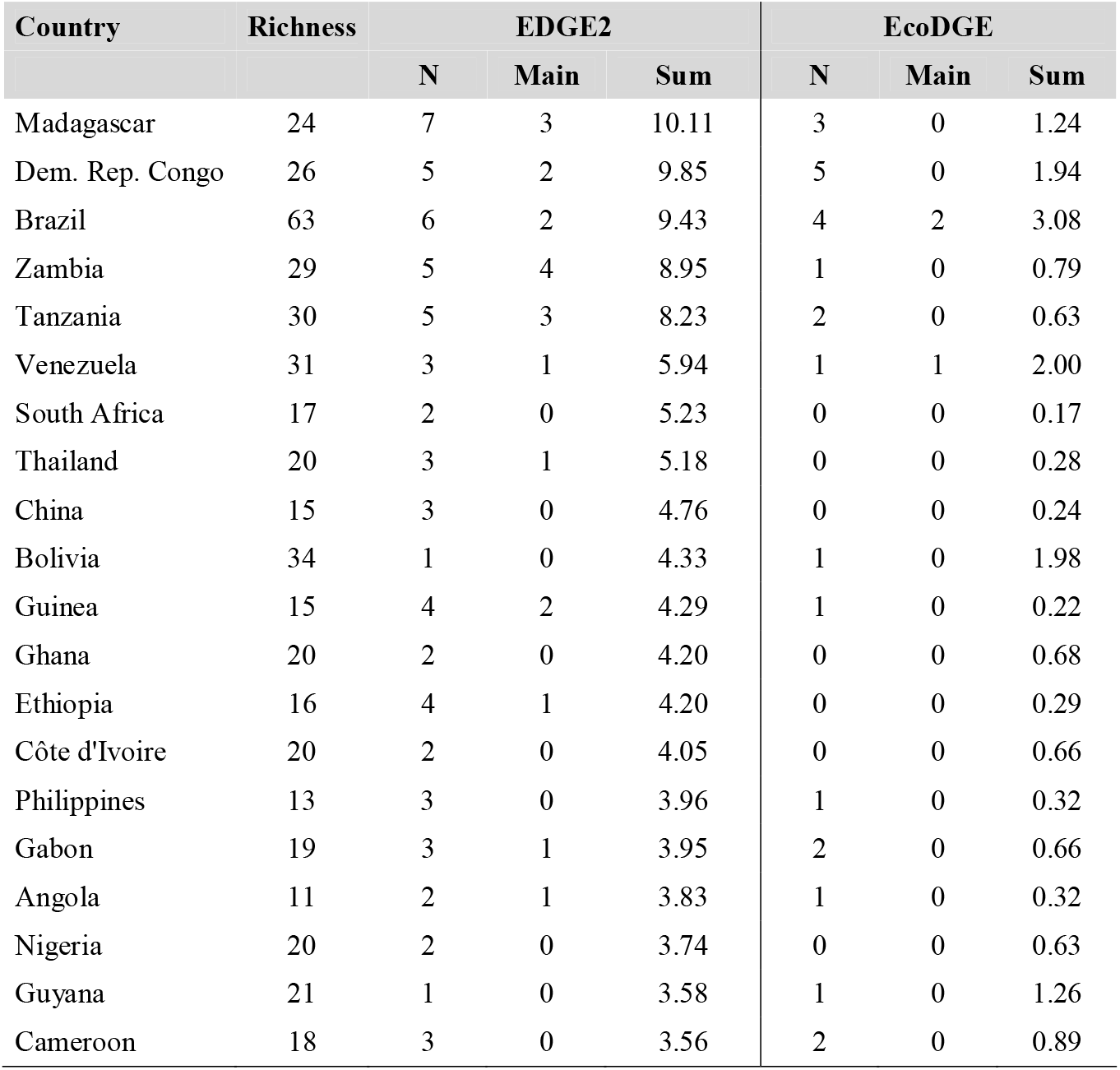
Top 20 countries in which *Scleria* is present ranked by their sum of EDGE2 scores. Richness: number of species present, N: number of EDGE2 and EcoDGE species, Main: number of species included in the main EDGE2 and EcoDGE lists, Sum: sum of EDGE2 and EcoDGE scores. Expanded results can be found in supplementary material Appendix 4.

**Table 5.**
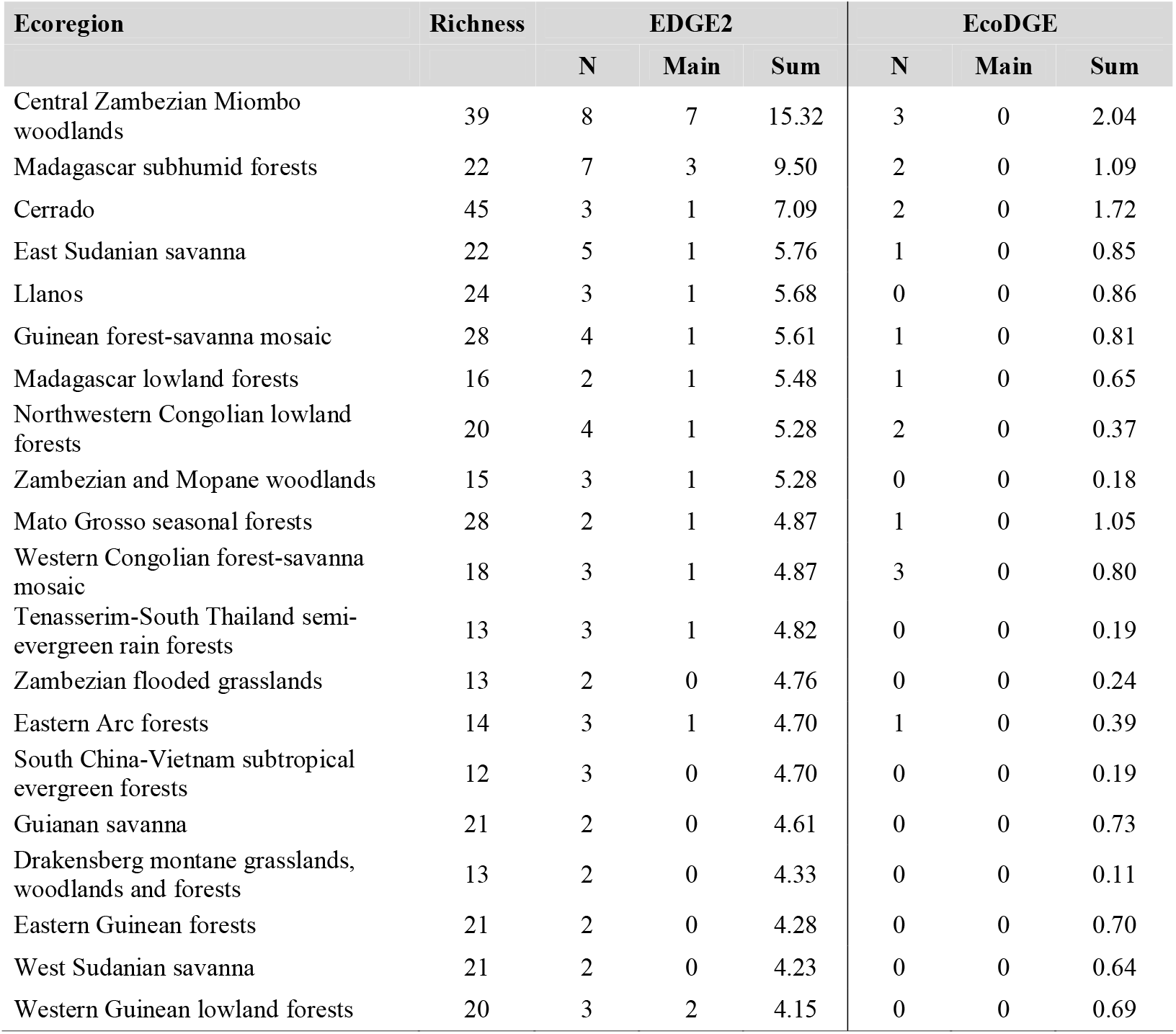
Top 20 ecoregions in which *Scleria* is present ranked by their sum of EDGE2 scores. Richness: number of species present, N: number of EDGE2 and EcoDGE species, Main: number of species included in the main EDGE2 and EcoDGE lists, Sum: sum of EDGE2 and EcoDGE scores. Expanded results can be found in supplementary material Appendix 5.

**Fig. 3.**
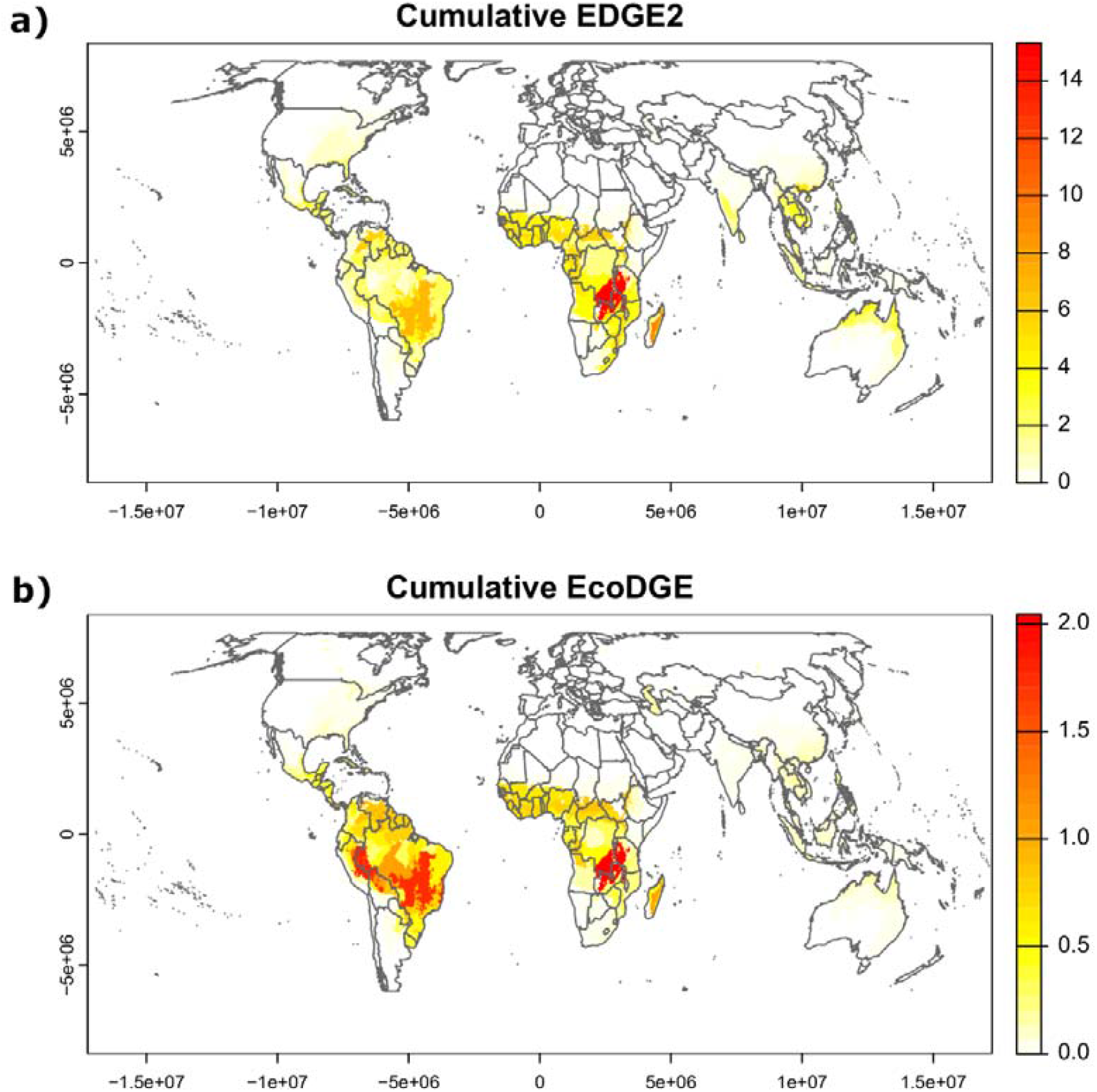
Ecoregions of remarkable interest for *Scleria* conservation according to their sum of EDGE2 scores (a) and sum of EcoDGE scores (b). The ecoregion definition follows Olson et al. (2001).

The countries with the highest cumulative EcoDGE scores were Brazil, Venezuela, Bolivia, Democratic Republic of the Congo, Peru and Colombia. The countries with the greatest number of listed EcoDGE species were Democratic Republic of the Congo, Brazil and Madagascar (Table 4). The ecoregions with the highest sum of EcoDGE scores and number of listed EcoDGE species were Central Zambezian Miombo woodlands, Southwest Amazon moist forests, Bahia coastal forests, Cerrado and Western Congolian forest-savanna mosaic (Table 5; Fig. 3a).

According to the assessment of the extent of remaining natural habitat and protected land in the world ecoregions by Dinerstein et al. (2017), there are 22 *Scleria* species which are restricted to ‘Nature Imperiled’ ecoregions (i.e., ecoregions where the percentage of natural habitat remaining and the amount of the total ecoregion that is protected is less than or equal to 20%) and 34 species only occur in ecoregions where ‘Nature Could Recover (i.e., ecoregions where the percentage of natural habitat remaining and area protected is less than 50% but more than 20% and that require restoration programs to reach the 50% of natural habitat).

## Discussion

In this study, we estimated the extinction risk for all species of the genus *Scleria*, as well as identified regions of special interest for its conservation. Our results suggest that half of *Scleria* species (49%, N=38) not yet included in the Red List are potentially threatened with extinction. Evolutionary and ecologically distinct and endangered *Scleria* mostly occur across African and South American regions.

### Preliminary assessments

Similar to previous studies, RF outperformed the method that explicitly followed Red List criterion B (Nic Lughadha et al. 2019) and distribution range properties were the best predictors for RF (Nic Lughadha et al. 2019; Bachman et al. 2023; Walker et al. 2023). Up to 15 species were classified as threatened by rCAT and non-threatened by RF; thus, RF set lower thresholds of extent of occurrence (EOO) to classify *Scleria* species as threatened. This seems a sensible approach to assess *Scleria* because the size of the distribution range as estimated from herbarium collections is potentially underestimated in most cases (see below), and widespread and locally dominant species are common (Holm et al. 1979; Galán Díaz et al. 2019).

Considering species included in the Red List and the results of RF, 26% of all *Scleria* species are potentially threatened with extinction. In a recent study, Bachman et al. (2023) obtained a similar estimate for Cyperaceae. They also found that, of the 21 families with more than 3,000 species, Cyperaceae showed the smallest proportion of threatened species. Whereas 18% of *Scleria* species currently included in the Red List are threatened, we found that 49% of unassessed species are potentially threatened with extinction. This could be because several Madagascan and South American *Scleria* species discovered over the last four decades were classified as threatened by RF, which might support that newly described species are geographically restricted and therefore less likely to be encountered in the wild (Brown et al. 2023). This is the case of *Scleria nusbaumeri, Scleria ankaratrensis, Scleria attenuatifolia, Scleria chasmema, Scleria millespicula, Scleria pernambucana, Scleria rubrostriata* and *Scleria tropicalis*. It is therefore necessary to prioritize the assessment of all unassessed species to get realistic estimates of extinction risk in the genus and to galvanize conservation support for these species.

Preliminary assessments are biased by the uncertainty and coverage of the taxonomic and geographical dimensions of the occurrence datasets (Meyer et al. 2016). We expect little uncertainty in the taxonomic dimension of our occurrence dataset because the records have been manually curated and homogenized using the latest list of accepted species names. Yet, there is a bias in the geographical coverage of *Scleria* collections, where the available resources for each country vary greatly and affect the density of observations as well as the number of specimens collected and digitized. For instance, the United States of America and Australia are the countries with the highest average number of *Scleria* observations per species with 276 and 273 respectively, five times more than the third and fourth countries in this ranking (Brazil and Mexico). This affects the assessments of species from less explored regions by potentially leading to an underestimation of the size of their distribution ranges and a subsequent overestimation of their extinction risk (Nic Lughadha et al. 2019).

### Evolutionarily distinct and globally endangered Scleria

Our analysis identified 23 species that met the criteria to be included in the EDGE2 main list. *Scleria porphyrocarpa* was the species with the highest EDGE2 and ED2 scores due to its high vulnerability (i.e., it was classified as threatened by both rCAT and RF) and high evolutionary distinctiveness. *Scleria porphyrocarpa* belongs to section *Corymbosae*, the oldest lineage in subgenus *Scleria* (crown age 22.28 Ma) which only includes four other species (Bauters et al. 2016, Larridon et al. 2021b). Section *Hypoporum* was represented in the EDGE2 main list by 17 species. This is due to two reasons: (i) molecular analyses indicated that section *Hypoporum* originated 8.4 Ma and showed a rapid increase in its diversification rate soon after (Larridon et al. 2021b) to become the richest section in *Scleria* (Bauters et al. 2019); (ii) it holds the highest percentage of threatened species among *Scleria* sections according to the Red List and our preliminary assessments (42%; Fig. 1b). This is important in the context of the new EDGE formulation, that accounts for the extinction risk of closely related species, because the deeper branches of a clade with a high proportion of threatened species will have a greater probability of being at risk, thus resulting in species belonging to it having higher EDGE scores (Gumbs et al. 2023). Finally, the three threatened species of section *Acriulus* were also included in the main list, as well as two Madagascan endemics from section *Abortivae*.

Almost half of *Scleria* species (129 species) are present in the 11 ecoregions with the greatest cumulative EDGE2 scores. Four African (Madagascar, D.R. Congo, Zambia and Tanzania) and two South American (Brazil, Venezuela) countries mainly covered these areas of special interest for conservation. Madagascar comprises 24 species of *Scleria* in two ecoregions (i.e., subhumid and lowland forests), including five threatened and two recently described species. Madagascan species belong to 11 sections out of the 17 sections recognized in *Scleria*, which highlights the importance of this country as a reservoir of *Scleria* diversity and the importance of long-distance dispersal in the genus (Galán Díaz et al. 2019). Four other ecoregions from the Afrotropics (i.e., Central Zambezian Miombo woodlands, East Sudanian Savanna, Guinean forest-savanna mosaic, and Northwestern Congolian lowland forests and Zambezian and Mopane woodlands; Olson et al. 2001) are identified as important conservation areas for *Scleria*. These ecoregions contain 67 species (26% species of *Scleria*, from 12 sections) of which 19 are endangered.

Brazil has the greatest *Scleria* richness (63 species) and ranks second in terms of cumulative EDGE2 scores. Two Brazilian ecoregions are shown as areas of special interest for the conservation of the evolutionary history of *Scleria*: Cerrado and Mato Grosso seasonal forests. These ecoregions include 49 species from 11 sections, of which 4 are endangered. The Venezuelan Llanos is another important ecoregion for evolutionary distinct and endangered *Scleria*. Unlike the Colombian Llanos, which also includes over 30 species, it includes species from old *Scleria* lineages (i.e., sections *Lithospermae* and *Foveolidia*; Larridon et al. 2021b) and two threatened species.

According to the revision of the world ecoregions of Dinerstein et al. (2017), which considered the World Database of Protected Areas (UNEP-WCMC and IUCN 2023) along with habitat assessments based on tree cover and human land use, two of the above-mentioned ecoregions (i.e., Madagascar subhumid forests and Guinean forest-savanna mosaic) are classified as ‘nature imperiled’ because the percentage of natural habitat remaining and protected area is less than 20%. Other ecoregions of interest for *Scleria* conservation that need active restoration programs to reach the 50% of protected natural habitat are Madagascar lowland forests, Central Zambezian Miombo woodlands and East Sudanian savanna.

### Ecologically distinct and globally endangered Scleria

We expect uncertainty in the identification of EcoDGE species because the results are largely dependent on the selection of traits, whereas the evolutionary relationship among *Scleria* species is well resolved at least at the section level (Bauters et al. 2016, 2018). In this study we used traits which are informative of species ecological strategies (Westoby 1998): plant longevity, leaf area, height and nutlet size. These traits indicate how plants cope with environmental nutrient stress and disturbances, the position of the species in the vertical light gradient of the vegetation, competitive vigor, dispersal distance and seed persistence in the soil bank (Pérez-Harguindeguy et al. 2013). We found ED2 and FUD were largely orthogonal; thus, EcoDGE based prioritizations yielded complementary results to the EDGE2 approach.

Our analyses identified six species in the main EcoDGE list. Unlike EDGE2, EcoDGE mostly points towards South American countries as reservoirs of distinctive and endangered species. This is because many EDGE2 species are phylogenetically restricted to section *Hypoporum* which has its center of diversity in tropical Africa, whereas EcoDGE species (i.e., those with great blade area and nutlet size) belong to different sections mostly restricted to South America. Eight South and Central American countries are ranked among the top 10 countries with the highest cumulative EcoDGE scores: Brazil, Venezuela, Bolivia, Peru, Colombia, Guyana, Dominican Republic and Costa Rica. This is due to the occurrence of stout species from three sections that are restricted or almost restricted to the Neotropics (Bauters et al. 2016): sections *Schizolepsis* (12 species), *Ophryoscleria* (11 species) and *Hymenolytrum* (13 species). Within these countries, the ecoregions that showed the greatest cumulative EcoDGE scores were Southwest Amazon moist forests, Bahia coastal forests, Cerrado and Iquitos Varzeá (87 species in total). The percentage of natural habitat remaining in these ecoregions is less than 50%, with restoration programs needed to exceed this threshold (Dinerstein et al. 2017). D. R. Congo and Madagascar were the African countries with the highest cumulative EcoDGE score.

Finally, it is worth mentioning that the identification of regions from Africa as areas of special interest for the conservation of *Scleria* reflects their great diversity as centers of diversification (Larridon et al. 2021b), but also our more comprehensive taxonomic knowledge of this group in particular countries. For instance, *Scleria* from Zambia, South Africa and Madagascar are better studied compared to other African countries thanks to the work of E.A. Robinson (Robinson 1966), E.F. Franklin Hennessy (Franklin Hennessy 1985) and H. Chermezon (Chermezon 1937). Thus, remarkable patterns of species richness in these countries might reflect different botanical collection efforts rather than true species accumulation.

### Conclusions

We provided the first global assessment of species and world regions of remarkable interest for *Scleria* conservation. We show that recent methodological advances in the identification of species at-risk of extinction in combination with herbarium data allow the implementation of the novel EDGE2 and EcoDGE frameworks in plant groups with well-resolved taxonomies. We found that half of *Scleria* species not yet included in the Red List are potentially at risk of extinction, and 21.5% of the species in the genus are restricted to ecoregions where the percentage of natural habitat is less than 50%. Phylogenetic and functional distinctiveness metrics were largely uncorrelated and EDGE2 and EcoDGE metrics point toward tropical regions of Africa and South America, respectively, as reservoirs of distinctive and endangered species.

## Acknowledgements

We thank Martin Xanthos from the Royal Botanic Gardens Kew, Germinal Rouhan and Thomas Haevermans from the Muséum National d’Histoire Naturelle and the Spanish Ecological Association of Terrestrial Ecology.

## Ethics approval

Not applicable.

## Consent to participate

Not applicable.

## Consent for publication

Not applicable.

## Availability of data and material (data transparency)

The data used to produce this study are publicly available at Zenodo (Galán Díaz et al. 2024).

## Code availability

The codes generated during the current study are available at GitHub (https://github.com/galanzse/Scleria_EDGE).

## Authors’ contributions

JGD and IL originally formulated the idea. JGD, HR and IL gathered and curated the data. JGD, SPB, FF and ME performed statistical analyses. JGD, SPB, FF, ME, HR and IL wrote the manuscript. JGD and IL acquired funding.

