## Supplementary Material for "Identifying conservation priorities of a pantropical plant lineage: a case study in Scleria (Cyperaceae)"

**Appendix 1.** Combined Red List results. This includes assessments published in the Red List of the IUCN version 2023-1 (RL IUCN), unpublished assessments (‘Assessed but not published’) and preliminary assessments (Threatened: T, Non-threatened: NT) of 67 species for which occurrence data was available. Preliminary assessments followed two approaches.

**(i) Random forest approach (RF).** We used as predictors: EOO, AOO, mean of latitudinal range, elevation, minimum human population density in 2020 (HPD; CIESIN 2018), human footprint index in 2013 (HFI; Venter et al. 2018), proportion of observations located in protected areas (UNEP-WCMC and IUCN 2023), annual mean temperature, minimum temperature of the coldest month, temperature annual range, annual precipitation, precipitation of the driest month and precipitation seasonality. We performed 10 repeats of 5-fold cross-validation to train and evaluate the model and retained 20% of data for external validation.

**(ii) rCAT.** We implemented the function ‘ConBatch’ from the R package ‘rCAT’ (Moat 2017) to assess species’ extinction risk following IUCN Criterion B based on EOO.

| **Scientific name** | **RL IUCN** | **RF** | **Rcat** | **Notes** |
| --- | --- | --- | --- | --- |
| *S. acanthocarpa* Boeckeler | LC |  |  | Included in RL 2023-1. |
| *S. achtenii* De Wild. | LC |  |  | Included in RL 2023-1. |
| *S. adpressohirta* (Kük.) E.A.Rob. | NE | T | T | Preliminary assessment. |
| *S. afroreflexa* Lye | EN |  |  | Included in RL 2023-1. |
| *S. alpina* Core | NE | NT | T | Preliminary assessment. |
| *S. amazonica* Camelb., M.T.Strong & Goetgh. | LC |  |  | Included in RL 2023-1. |
| *S. anceps* Liebm. | NE | T | T | Preliminary assessment. |
| *S. andringitrensis* Cherm. | EN |  |  | Included in RL 2023-1. |
| *S. angusta* Nees ex Kunth | LC |  |  | Included in RL 2023-1. |
| *S. angustifolia* E.A.Rob. | LC |  |  | Included in RL 2023-1. |
| *S. ankaratrensis* Bauters | DD | T | T | Included in RL 2023-1. |
| *S. annularis* Steud. | LC |  |  | Assessed but not published. |
| *S. anomala* (Steud.) J.Raynal | NE | T | T | Preliminary assessment. |
| *S. arcuata* E.A.Rob. | NE | T | T | Preliminary assessment. |
| *S. arenaria* T.Koyama | NE | T | T | Preliminary assessment. |
| *S. arguta* (Nees) Steud. | LC |  |  | Included in RL 2023-1. |
| *S. aromatica* Core | NE | NT | NT | Preliminary assessment. |
| *S. atroglumis* D.A.Simpson | LC |  |  | Included in RL 2023-1. |
| *S. attenuatifolia* M.T.Strong | NE | T | T | Preliminary assessment. |
| *S. aurantiaca* Lye | CR |  |  | Included in RL 2023-1. |
| *S. aureovillosa* Kiaos. & K.Wangwasit | NE | T | T | Preliminary assessment. |
| *S. balansae* Maury ex Micheli | LC |  |  | Included in RL 2023-1. |
| *S. baldwinii* (Torr.) Steud. | LC |  |  | Included in RL 2023-1. |
| *S. bambariensis* Cherm. | LC |  |  | Assessed but not published. |
| *S. baroni-clarkei* De Wild. | EN |  |  | Included in RL 2023-1. |
| *S. baronii* C.B.Clarke ex Cherm. | LC |  |  | Included in RL 2023-1. |
| *S. bellii* LeBlond | NE | NT | NT | Preliminary assessment. |
| *S. benthamii* C.B.Clarke | NE | NT | T | Preliminary assessment. |
| *S. bequaertii* De Wild. | LC |  |  | Included in RL 2023-1. |
| *S. biflora* Roxb. | LC |  |  | Included in RL 2023-1. |
| *S. boivinii* Steud. | LC |  |  | Included in RL 2023-1. |
| *S. boniana* Boeckeler | NE | NT | T | Preliminary assessment. |
| *S. borii* D.M.Verma | NE | NT | T | Preliminary assessment. |
| *S.* *bourgeaui* Boeckeler | LC |  |  | Included in RL 2023-1. |
| *S. bracteata* Cav. | LC |  |  | Included in RL 2023-1. |
| *S. bradei* Gross | NE | T | T | Preliminary assessment. |
| *S. brownii* Kunth | LC |  |  | Included in RL 2023-1. |
| *S. bulbifera* Hochst. ex A.Rich. | LC |  |  | Included in RL 2023-1. |
| *S. burchellii* C.B.Clarke | EN |  |  | Included in RL 2023-1. |
| *S. calcicola* E.A.Rob. | EN |  |  | Assessed but not published. |
| *S. camaratensis* Core | NE | NT | NT | Preliminary assessment. |
| *S. canescens* Boeckeler | LC |  |  | Assessed but not published. |
| *S. carphiformis* Ridl. | LC |  |  | Included in RL 2023-1. |
| *S. castanea* Core | LC |  |  | Included in RL 2023-1. |
| *S. catophylla* C.B.Clarke | LC |  |  | Included in RL 2023-1. |
| *S. chasmema* Bonet Mayedo & W.W.Thomas | NE | T | T | Preliminary assessment. |
| *S. cheekii* Bauters | VU |  |  | Included in RL 2023-1. |
| *S. chevalieri* J.Raynal | EX |  |  | Included in RL 2023-1. |
| *S. chlorantha* Boeckeler | NE | T | T | Preliminary assessment. |
| *S. chlorocalyx* E.A.Rob. | NE | NT | T | Preliminary assessment. |
| *S. ciliaris* Nees | LC |  |  | Included in RL 2023-1. |
| *S. ciliata* Michx. | LC |  |  | Included in RL 2023-1. |
| *S. clarkei* Lindm. | NE | T | T | Preliminary assessment. |
| *S. clathrata* Hochst. ex A.Rich. | NE | NT | NT | Preliminary assessment. |
| *S. colorata* Core | NE | NT | T | Preliminary assessment. |
| *S. comosa* (Nees) Steud. | LC |  |  | Included in RL 2023-1. |
| *S. composita* (Nees) Boeckeler | LC |  |  | Included in RL 2023-1. |
| *S. corymbosa* Roxb. | LC |  |  | Assessed but not published. |
| *S. cuyabensis* Pilg. | LC |  |  | Included in RL 2023-1. |
| *S. cyathophora* Holttum | NE | NT | T | Preliminary assessment. |
| *S. cyperina* Kunth | LC |  |  | Included in RL 2023-1. |
| *S. delicatula* Nelmes | NT |  |  | Included in RL 2023-1. |
| *S. densispicata* (C.B.Clarke) J.Kern | NE | T | T | Preliminary assessment. |
| *S. depressa* (C.B.Clarke) Nelmes | LC |  |  | Included in RL 2023-1. |
| *S. didina* Bonet Mayedo & W.W.Thomas | NE | NT | NT | Preliminary assessment. |
| *S. distans* Poir. | LC |  |  | Included in RL 2023-1. |
| *S. dregeana* Kunth | LC |  |  | Included in RL 2023-1. |
| *S. dulungensis* P.C.Li | NE | NT | T | Preliminary assessment. |
| *S. eggersiana* Boeckeler | LC |  |  | Included in RL 2023-1. |
| *S. erythrorrhiza* Ridl. | LC |  |  | Included in RL 2023-1. |
| *S. filiculmis* Boeckeler | NE | T | T | Preliminary assessment. |
| *S. flagellum-nigrorum* P.J.Bergius | LC |  |  | Included in RL 2023-1. |
| *S. flexuosa* Boeckeler | LC |  |  | Included in RL 2023-1. |
| *S. foliosa* Hochst. ex A.Rich. | LC |  |  | Included in RL 2023-1. |
| *S. foveolata* Cav. | NE | T | T | Preliminary assessment. |
| *S. fulvipilosa* E.A.Rob. | EN |  |  | Included in RL 2023-1. |
| *S. gaertneri* Raddi | LC |  |  | Included in RL 2023-1. |
| *S. georgiana* Core | LC |  |  | Included in RL 2023-1. |
| *S. glabra* Boeckeler | LC |  |  | Included in RL 2023-1. |
| *S. globonux* C.B.Clarke | LC |  |  | Included in RL 2023-1. |
| *S. glomerulata* Oliv. | CR |  |  | Included in RL 2023-1. |
| *S. goossensii* De Wild. | LC |  |  | Included in RL 2023-1. |
| *S. gracillima* Boeckeler | LC |  |  | Included in RL 2023-1. |
| *S. greigiifolia* (Ridl.) C.B.Clarke | LC |  |  | Included in RL 2023-1. |
| *S. guineensis* J.Raynal | CR |  |  | Included in RL 2023-1. |
| *S. harlandii* Hance | LC |  |  | Included in RL 2023-1. |
| *S. havanensis* Britton | LC |  |  | Included in RL 2023-1. |
| *S. hildebrandtii* Boeckeler | EN |  |  | Included in RL 2023-1. |
| *S. hilsenbergii* Ridl. | LC |  |  | Included in RL 2023-1. |
| *S. hirtella* Sw. | LC |  |  | Included in RL 2023-1. |
| *S. hispidior* (C.B.Clarke) Nelmes | LC |  |  | Included in RL 2023-1. |
| *S. hispidula* Hochst. ex A.Rich. | LC |  |  | Included in RL 2023-1. |
| *S. huberi* C.B.Clarke | LC |  |  | Assessed but not published. |
| *S. indica* D.M.Verma & Veena Chandra | NE | NT | T | Preliminary assessment. |
| *S. induta* Turrill | NE | T | NT | Preliminary assessment. |
| *S. interrupta* Rich. | LC |  |  | Included in RL 2023-1. |
| *S. iostephana* Nelmes | LC |  |  | Included in RL 2023-1. |
| *S. junghuhniana* Boeckeler | NE | T | T | Preliminary assessment. |
| *S. kerrii* Turrill | LC |  |  | Included in RL 2023-1. |
| *S. khasiana* Boeckeler | NE | T | T | Preliminary assessment. |
| *S. lacustris* C.Wright | LC |  |  | Included in RL 2023-1. |
| *S. lagoensis* Boeckeler | LC |  |  | Included in RL 2023-1. |
| *S. latifolia* Sw. | LC |  |  | Included in RL 2023-1. |
| *S. laxa* R.Br. | LC |  |  | Included in RL 2023-1. |
| *S. laxiflora* Gross | LC |  |  | Included in RL 2023-1. |
| *S. leptostachya* Kunth | LC |  |  | Included in RL 2023-1. |
| *S. levis* Retz. | LC |  |  | Assessed but not published. |
| *S. liberica* Bauters | VU |  |  | Included in RL 2023-1. |
| *S. lingulata* C.B.Clarke | LC |  |  | Assessed but not published. |
| *S. lithosperma* (L.) Sw. | LC |  |  | Included in RL 2023-1. |
| *S. longispiculata* Nelmes | LC |  |  | Included in RL 2023-1. |
| *S. lucentinigricans* E.A.Rob. | NE | NT | T | Preliminary assessment. |
| *S. macbrideana* Gross | LC |  |  | Included in RL 2023-1. |
| *S. mackaviensis* Boeckeler | LC |  |  | Included in RL 2023-1. |
| *S. macrogyne* C.B.Clarke | LC |  |  | Included in RL 2023-1. |
| *S. macrophylla* J.Presl & C.Presl | LC |  |  | Included in RL 2023-1. |
| *S. madagascariensis* Boeckeler | EN |  |  | Included in RL 2023-1. |
| *S. martii* (Nees) Steud. | LC |  |  | Assessed but not published. |
| *S. maypurensis* Bauters | CR |  |  | Included in RL 2023-1. |
| *S. melanomphala* Kunth | LC |  |  | Included in RL 2023-1. |
| *S. melanotricha* Hochst. & A.Rich. | LC |  |  | Included in RL 2023-1. |
| *S. microcarpa* Nees ex Kunth | LC |  |  | Included in RL 2023-1. |
| *S.* *mikawana* Makino | LC |  |  | Included in RL 2023-1. |
| *S. millespicula* T.Koyama | NE | T | T | Preliminary assessment. |
| *S. minor* (Britton) W.Stone | LC |  |  | Included in RL 2023-1. |
| *S. mitis* P.J.Bergius | LC |  |  | Included in RL 2023-1. |
| *S. mongomoensis* Bauters | EN |  |  | Included in RL 2023-1. |
| *S. monticola* Nelmes ex Napper | NE | NT | T | Preliminary assessment. |
| *S. motleyi* C.B.Clarke | NE | NT | NT | Preliminary assessment. |
| *S. mucronata* Poir. | LC |  |  | Included in RL 2023-1. |
| *S. muehlenbergii* Steud. | LC |  |  | Included in RL 2023-1. |
| *S. multilacunosa* T.Koyama | NE | T | T | Preliminary assessment. |
| *S. myricocarpa* Kunth | LC |  |  | Assessed but not published. |
| *S. natalensis* Boeckeler ex C.B.Clarke | LC |  |  | Included in RL 2023-1. |
| *S. naumanniana* Boeckeler | LC |  |  | Included in RL 2023-1. |
| *S. neesii* Kunth | LC |  |  | Included in RL 2023-1. |
| *S. neocaledonica* Rendle | NE | T | T | Preliminary assessment. |
| *S. neogranatensis* C.B.Clarke | LC |  |  | Included in RL 2023-1. |
| *S. novae-hollandiae* Boeckeler | LC |  |  | Assessed but not published. |
| *S. nusbaumeri* Bauters | NE | T | T | Preliminary assessment. |
| *S. nyasensis* C.B.Clarke | LC |  |  | Included in RL 2023-1. |
| *S. oblata* S.T.Blake ex J.Kern | LC |  |  | Included in RL 2023-1. |
| *S. obtusa* Core | LC |  |  | Assessed but not published. |
| *S. oligantha* Michx. | LC |  |  | Included in RL 2023-1. |
| *S. oligochondra* Nelmes | NE | T | T | Preliminary assessment. |
| *S. orchardii* C.D.Adams | CR |  |  | Included in RL 2023-1. |
| *S. ovinux* J.Raynal ex Fosberg | NE | T | T | Preliminary assessment. |
| *S. pachyrrhyncha* Nelmes | EN |  |  | Included in RL 2023-1. |
| *S. panicoides* Kunth | LC |  |  | Included in RL 2023-1. |
| *S. pantadenia* Meganck & Bauters | VU |  |  | Included in RL 2023-1. |
| *S. parallella* C.B.Clarke | NE | NT | NT | Preliminary assessment. |
| *S. parvula* Steud. | LC |  |  | Included in RL 2023-1. |
| *S. patula* E.A.Rob. | NE | T | T | Preliminary assessment. |
| *S. pauciflora* Muhl. ex Willd. | LC |  |  | Included in RL 2023-1. |
| *S. paupercula* E.A.Rob. | LC |  |  | Assessed but not published. |
| *S. pedicellata* Bauters | EN |  |  | Included in RL 2023-1. |
| *S. pergracilis* (Nees) Kunth | LC |  |  | Included in RL 2023-1. |
| *S. pernambucana* Luceño & M.Alves | NE | T | T | Preliminary assessment. |
| *S. perpusilla* Cherm. | EN |  |  | Included in RL 2023-1. |
| *S. pilosa* Boeckeler | NE | T | T | Preliminary assessment. |
| *S. pilosissima* Britton | NE | NT | T | Preliminary assessment. |
| *S. plusiophylla* Steud. | LC |  |  | Included in RL 2023-1. |
| *S. poeppigii* (Nees) Steud. | NE | T | T | Preliminary assessment. |
| *S. poiformis* Retz. | LC |  |  | Included in RL 2023-1. |
| *S. polycarpa* Boeckeler | LC |  |  | Included in RL 2023-1. |
| *S. polyrrhiza* E.A.Rob. | EN |  |  | Included in RL 2023-1. |
| *S. pooides* Ridl. | LC |  |  | Included in RL 2023-1. |
| *S. porphyrocarpa* E.A.Rob. | NE | T | T | Preliminary assessment. |
| *S. procumbens* E.A.Rob. | NE | NT | T | Preliminary assessment. |
| *S. pseudohispidior* Bauters | VU |  |  | Included in RL 2023-1. |
| *S. psilorrhiza* C.B.Clarke | LC |  |  | Assessed but not published. |
| *S. pulchella* Ridl. | CR |  |  | Included in RL 2023-1. |
| *S. purdiei* C.B.Clarke | LC |  |  | Included in RL 2023-1. |
| *S. purpurascens* Steud. | LC |  |  | Assessed but not published. |
| *S. pusilla* Pilg. | LC |  |  | Included in RL 2023-1. |
| *S. racemosa* Poir. | LC |  |  | Included in RL 2023-1. |
| *S. radula* Hance | LC |  |  | Assessed but not published. |
| *S. ramosa* C.B.Clarke | LC |  |  | Included in RL 2023-1. |
| *S. rehmannii* C.B.Clarke | LC |  |  | Included in RL 2023-1. |
| *S. remota* Ridl. | NE | T | T | Preliminary assessment. |
| *S. reticularis* Michx. | LC |  |  | Included in RL 2023-1. |
| *S. richardsiae* E.A.Rob. | EN |  |  | Included in RL 2023-1. |
| *S. robinsoniana* J.Raynal | NT |  |  | Included in RL 2023-1. |
| *S. robusta* Camelb. & Goetgh. | LC |  |  | Included in RL 2023-1. |
| *S. rosea* Cherm. | LC |  |  | Included in RL 2023-1. |
| *S. rubrostriata* A.C.Araújo & N.A.Brummitt | NE | T | T | Preliminary assessment. |
| *S. rugosa* R.Br. | LC |  |  | Included in RL 2023-1. |
| *S. rutenbergiana* Boeckeler | NT |  |  | Included in RL 2023-1. |
| *S. scabra* Willd. | LC |  |  | Included in RL 2023-1. |
| *S. scabriuscula* Schltdl. | NE | NT | NT | Preliminary assessment. |
| *S. schiedeana* Schltdl. | LC |  |  | Included in RL 2023-1. |
| *S. schimperiana* Boeckeler | LC |  |  | Included in RL 2023-1. |
| *S. schulzii* Barros | NE | NT | T | Preliminary assessment. |
| *S. scrobiculata* Nees & Meyen | LC |  |  | Assessed but not published. |
| *S. secans* (L.) Urb. | LC |  |  | Included in RL 2023-1. |
| *S. sellowiana* Kunth | LC |  |  | Assessed but not published. |
| *S. setulosociliata* Boeckeler | LC |  |  | Included in RL 2023-1. |
| *S. sheilae* J.Raynal | CR |  |  | Included in RL 2023-1. |
| *S. sieberi* Nees ex Kunth | NE | NT | T | Preliminary assessment. |
| *S. skutchii* M.T.Strong & J.R.Grant | LC |  |  | Included in RL 2023-1. |
| *S. sobolifera* E.F.Franklin | LC |  |  | Included in RL 2023-1. |
| *S. sororia* Kunth | LC |  |  | Included in RL 2023-1. |
| *S. sphacelata* F.Muell. | LC |  |  | Included in RL 2023-1. |
| *S. spicata* (Spreng.) J.F.Macbr. | VU |  |  | Included in RL 2023-1. |
| *S. spiciformis* Benth. | LC |  |  | Included in RL 2023-1. |
| *S. splitgerberiana* Henrard ex Uittien | LC |  |  | Included in RL 2023-1. |
| *S. sprucei* C.B.Clarke | LC |  |  | Included in RL 2023-1. |
| *S. staheliana* Uittien | LC |  |  | Included in RL 2023-1. |
| *S. stipitata* Uittien | NE | NT | NT | Preliminary assessment. |
| *S. stipularis* Nees | LC |  |  | Included in RL 2023-1. |
| *S. stocksiana* Boeckeler | NE | NT | T | Preliminary assessment. |
| *S. suaveolens* Nelmes | LC |  |  | Included in RL 2023-1. |
| *S. suffulta* C.B.Clarke | NE | T | T | Preliminary assessment. |
| *S. sumatrensis* Retz. | LC |  |  | Included in RL 2023-1. |
| *S. tenacissima* (Nees) Steud. | LC |  |  | Assessed but not published. |
| *S. tenella* Kunth | LC |  |  | Included in RL 2023-1. |
| *S. tepuiensis* Core | LC |  |  | Included in RL 2023-1. |
| *S. terrestris* (L.) Fassett | LC |  |  | Included in RL 2023-1. |
| *S. tessellata* Willd. | LC |  |  | Included in RL 2023-1. |
| *S. testacea* Nees ex Kunth | LC |  |  | Included in RL 2023-1. |
| *S. thwaitesiana* Boeckeler | NE | T | NT | Preliminary assessment. |
| *S. tonkinensis* C.B.Clarke | LC |  |  | Included in RL 2023-1. |
| *S. transvaalensis* E.F.Franklin | NT |  |  | Included in RL 2023-1. |
| *S. trialata* Poir. | LC |  |  | Included in RL 2023-1. |
| *S. tricholepis* Nelmes | VU |  |  | Included in RL 2023-1. |
| *S. tricristata* Meganck & Bauters | EN |  |  | Assessed but not published. |
| *S. tricuspidata* S.T.Blake | LC |  |  | Assessed but not published. |
| *S. triglomerata* Michx. | LC |  |  | Included in RL 2023-1. |
| *S. triquetra* M.T.Strong | LC |  |  | Assessed but not published. |
| *S. tropicalis* M.T.Strong | NE | T | T | Preliminary assessment. |
| *S. tryonii* Domin | NE | T | T | Preliminary assessment. |
| *S. uleana* Boeckeler | NE | NT | NT | Preliminary assessment. |
| *S. unguiculata* E.A.Rob. | LC |  |  | Included in RL 2023-1. |
| *S. vaginata* Steud. | LC |  |  | Assessed but not published. |
| *S. variegata* (Nees) Steud. | LC |  |  | Assessed but not published. |
| S. *venezuelensis* Core | NE | T | T | Preliminary assessment. |
| *S. verrucosa* Willd. | LC |  |  | Included in RL 2023-1. |
| *S. verticillata* Muhl. ex Willd. | LC |  |  | Included in RL 2023-1. |
| *S. veseyfitzgeraldii* E.A.Rob. | LC |  |  | Assessed but not published. |
| *S. violacea* Pilg. | NE | NT | NT | Preliminary assessment. |
| S*. virgata* (Nees) Steud. | LC |  |  | Included in RL 2023-1. |
| *S. vogelii* C.B.Clarke | LC |  |  | Included in RL 2023-1. |
| *S. warmingiana* Boeckeler | NE | T | T | Preliminary assessment. |
| *S. welwitschii* C.B.Clarke | LC |  |  | Included in RL 2023-1. |
| *S. williamsii* Gross | EN |  |  | Assessed but not published. |
| *S. woodii* C.B.Clarke | LC |  |  | Included in RL 2023-1. |
| *S. wrightiana* Boeckeler | NE | NT | NT | Preliminary assessment. |
| *S. xerophila* E.A.Rob. | NE | T | T | Preliminary assessment. |
| *S. zambesica* E.A.Rob. | NE | T | T | Preliminary assessment. |

Species without occurrence data: *S. assamica* (C.B.Clarke) D.M.Verma, *S. depauperata* Boeckeler, *S. elongatissima* Piérart, *S. hirta* Boeckeler, *S. jiangchengensis* Y.Y.Qian, *S. lateritica* Nelmes, *S. macrolomioides* H.Pfeiff., *S. mutoensis* Nakai, *S. papuana* J.Kern, *S. scandens* Core, *S. schenckiana* Boeckeler and *S. swamyi* Govind.

**Appendix 2.** *Scleria* EDGE2 borderline and watch lists (Gumbs et al., 2022). Borderline list: threatened species whose distribution of EDGE2 scores rank above the median of the entire genus at least 80% of the time. Watch list: nonthreatened species whose distribution of EDGE2 scores rank above the median of the entire genus at least 95% of the times. EDGE2: Evolutionarily Distinct and Globally Endangered metric, ED2: evolutionary distinctiveness.

| **List** | **Scientific name** | **Section** | **EDGE2 (mean** ± **sd)** | **ED2**  **(mean** ± **sd)** |
| --- | --- | --- | --- | --- |
| Borderline | *S. chasmema* Bonet Mayedo & W.W.Thomas | *Virgatae* | 0.76 ± 1.85 | 1.57 ± 2.53 |
|  | *S. guineensis* J.Raynal | *Hypoporum* | 0.76 ± 0.85 | 0.85 ± 0.9 |
|  | *S. glomerulata* Oliv. | *Hypoporum* | 0.74 ± 0.85 | 0.85 ± 0.9 |
|  | *S. sheilae* J.Raynal | *Hypoporum* | 0.74 ± 0.81 | 0.81 ± 0.88 |
|  | *S. baroni-clarkei* De Wild. | *Foveolidia* | 0.59 ± 0.33 | 1.34 ± 0.6 |
|  | *S. poeppigii* (Nees) Steud. | *Hymenolytrum* | 0.58 ± 1.16 | 1.16 ± 1.58 |
|  | *S. densispicata* (C.B.Clarke) J.Kern | *Browniae* | 0.45 ± 3.04 | 0.88 ± 5.1 |
|  | *S. bradei* Gross | *Hypoporum* | 0.43 ± 0.77 | 0.83 ± 0.99 |
|  | *S. neocaledonica* Rendle | *Browniae* | 0.42 ± 3.61 | 0.9 ± 5.34 |
|  | *S. andringitrensis* Cherm. | *Hypoporum* | 0.42 ± 0.45 | 0.86 ± 0.87 |
|  | *S. oligochondra* Nelmes | *Ophryoscleria* | 0.41 ± 0.94 | 0.91 ± 1.39 |
|  | *S. perpusilla* Cherm. | *Hypoporum* | 0.41 ± 0.47 | 0.87 ± 0.93 |
|  | *S. afroreflexa* Lye | *Hypoporum* | 0.4 ± 0.54 | 0.86 ± 1.02 |
|  | *S. chlorantha* Boeckeler | *Naumannianae* | 0.4 ± 1.68 | 0.86 ± 2.42 |
|  | *S. remota* Ridl. | *Hypoporum* | 0.39 ± 0.67 | 0.77 ± 0.94 |
|  | *S. filiculmis* Boeckeler | *Hypoporum* | 0.38 ± 0.59 | 0.89 ± 0.92 |
|  | *S. tropicalis* M.T.Strong | *Ophryoscleria* | 0.37 ± 0.91 | 0.72 ± 1.29 |
| Watch | *S. transvaalensis* E.F.Franklin | *Acriulus* | 1.52 ± 0.57 | 13.88 ± 4.08 |
|  | *S. corymbosa* Roxb. | *Corymbosae* | 1.29 ± 0.59 | 22.28 ± 2.18 |
|  | *S. lithosperma* (L.) Sw. | *Lithospermae* | 1.22 ± 0.52 | 20.89 ± 0 |
|  | *S. melanomphala* Kunth | *Melanomphalae* | 1.15 ± 0.5 | 19.04 ± 0 |
|  | *S. tonkinensis* C.B.Clarke | *Corymbosae* | 0.82 ± 0.36 | 13.91 ± 1.63 |

**Appendix 3.** *Scleria* EcoDGE borderline list (Griffith et al. 2022; Gumbs et al. 2023): threatened species whose distribution of EcoDGE scores rank above the median of the entire genus at least 80% of the time. EcoDGE: Ecologically Distinct and Globally Endangered metric, FUD: functional distinctiveness.

| **List** | **Scientific name** | **Section** | **EcoDGE (mean** ± **sd)** | **FUD**  **(mean** ± **sd)** |
| --- | --- | --- | --- | --- |
|  | *S. porphyrocarpa* E.A.Rob. | *Corymbosae* | 1.14 ± 1.45 | 2.05 ± 2.1 |
|  | *S. williamsii* Gross | *Scleria* | 0.86 ± 1.33 | 1.83 ± 2.56 |
|  | *S. chlorantha* Boeckeler | *Naumannianae* | 0.84 ± 1.29 | 1.8 ± 1.78 |
|  | *S. attenuatifolia* M.T.Strong | *Schizolepis* | 0.54 ± 1.01 | 1.13 ± 1.36 |
|  | *S. foveolata* Cav. | *Schizolepis* | 0.46 ± 1.44 | 0.75 ± 2.07 |
|  | *S. khasiana* Boeckeler | *Elatae* | 0.31 ± 0.47 | 0.62 ± 0.63 |
|  | *S. madagascariensis* Boeckeler | *Abortivae* | 0.24 ± 0.15 | 0.49 ± 0.28 |
|  | *S. bradei* Gross | *Hypoporum* | 0.23 ± 0.4 | 0.67 ± 0.55 |
|  | *S. induta* Turrill | *Acriulus* | 0.2 ± 0.3 | 0.63 ± 0.38 |
|  | *S. nusbaumeri* Bauters | *Abortivae* | 0.2 ± 0.25 | 0.42 ± 0.35 |
| Borderline | *S. aurantiaca* Lye | *Foveolidia* | 0.18 ± 0.15 | 0.2 ± 0.16 |
|  | *S. warmingiana* Boeckeler | *Scleria* | 0.17 ± 0.26 | 0.54 ± 0.35 |
|  | *S. burchellii* C.B.Clarke | *Hypoporum* | 0.15 ± 0.12 | 0.28 ± 0.24 |
|  | *S. densispicata* (C.B.Clarke) J.Kern | *Browniae* | 0.13 ± 0.15 | 0.27 ± 0.2 |
|  | *S. pachyrrhyncha* Nelmes | *Acriulus* | 0.12 ± 0.07 | 0.27 ± 0.14 |
|  | *S. hildebrandtii* Boeckeler | *Foveolidia* | 0.12 ± 0.07 | 0.21 ± 0.13 |
|  | *S. adpressohirta* (KÃ¼k.) E.A.Rob. | *Foveolidia* | 0.11 ± 0.2 | 0.18 ± 0.27 |
|  | *S. arcuata* E.A.Rob. | *Foveolidia* | 0.1 ± 0.3 | 0.2 ± 0.42 |
|  | *S. suffulta* C.B.Clarke | *Elatae* | 0.1 ± 0.12 | 0.21 ± 0.16 |
|  | *S. glomerulata* Oliv. | *Hypoporum* | 0.1 ± 0.05 | 0.1 ± 0.05 |
|  | *S. oligochondra* Nelmes | *Ophryoscleria* | 0.1 ± 0.14 | 0.18 ± 0.21 |
|  | *S. patula* E.A.Rob. | *Foveolidia* | 0.09 ± 0.08 | 0.2 ± 0.09 |
|  | *S. pulchella* Ridl. | *Hypoporum* | 0.08 ± 0.06 | 0.09 ± 0.06 |
|  | *S. neocaledonica* Rendle | *Browniae* | 0.07 ± 0.1 | 0.13 ± 0.13 |
|  | *S. anomala* (Steud.) J.Raynal | *Elatae* | 0.06 ± 0.21 | 0.11 ± 0.29 |
|  | *S. guineensis* J.Raynal | *Hypoporum* | 0.06 ± 0.04 | 0.06 ± 0.04 |
|  | *S. cheekii* Bauters | *Hypoporum* | 0.06 ± 0.04 | 0.24 ± 0.17 |
|  | *S. aureovillosa* Kiaos. & K.Wangwasit | *Foveolidia* | 0.05 ± 0.05 | 0.11 ± 0.06 |
|  | *S. baroni-clarkei* De Wild. | *Foveolidia* | 0.05 ± 0.12 | 0.1 ± 0.24 |
|  | *S. multilacunosa* T.Koyama | *Foveolidia* | 0.04 ± 0.04 | 0.08 ± 0.05 |
|  | *S. mongomoensis* Bauters | *Hypoporum* | 0.04 ± 0.07 | 0.07 ± 0.14 |
|  | *S. sheilae* J.Raynal | *Hypoporum* | 0.03 ± 0.01 | 0.03 ± 0.01 |

**Appendix 4.** Countries in which *Scleria* is present ranked by their sum of EDGE2 scores. Richness: number of species present, N: number of listed EDGE2 and EcoDGE species, Main: number of species included in the main EDGE2 and EcoDGE lists, Sum: sum of EDGE2 and EcoDGE scores.

| **Country** | **Richness** | **EDGE2** | | | **EcoDGE** | | |
| --- | --- | --- | --- | --- | --- | --- | --- |
|  |  | **N** | **Main** | **Sum** | **N** | **Main** | **Sum** |
| Madagascar | 24 | 7 | 3 | 10.11 | 3 | 0 | 1.24 |
| Dem. Rep. Congo | 26 | 5 | 2 | 9.85 | 5 | 0 | 1.94 |
| Brazil | 63 | 6 | 2 | 9.43 | 4 | 2 | 3.08 |
| Zambia | 29 | 5 | 4 | 8.95 | 1 | 0 | 0.79 |
| Tanzania | 30 | 5 | 3 | 8.23 | 2 | 0 | 0.63 |
| Venezuela | 31 | 3 | 1 | 5.94 | 1 | 1 | 2.00 |
| South Africa | 17 | 2 | 0 | 5.23 | 0 | 0 | 0.17 |
| Thailand | 20 | 3 | 1 | 5.18 | 0 | 0 | 0.28 |
| China | 15 | 3 | 0 | 4.76 | 0 | 0 | 0.24 |
| Bolivia | 34 | 1 | 0 | 4.33 | 1 | 0 | 1.98 |
| Guinea | 15 | 4 | 2 | 4.29 | 1 | 0 | 0.22 |
| Ghana | 20 | 2 | 0 | 4.20 | 0 | 0 | 0.68 |
| Ethiopia | 16 | 4 | 1 | 4.20 | 0 | 0 | 0.29 |
| Côte d'Ivoire | 20 | 2 | 0 | 4.05 | 0 | 0 | 0.66 |
| Philippines | 13 | 3 | 0 | 3.96 | 1 | 0 | 0.32 |
| Gabon | 19 | 3 | 1 | 3.95 | 2 | 0 | 0.66 |
| Angola | 11 | 2 | 1 | 3.83 | 1 | 0 | 0.32 |
| Nigeria | 20 | 2 | 0 | 3.74 | 0 | 0 | 0.63 |
| Guyana | 21 | 1 | 0 | 3.58 | 1 | 0 | 1.26 |
| Cameroon | 18 | 3 | 0 | 3.56 | 2 | 0 | 0.89 |
| Australia | 22 | 1 | 0 | 3.51 | 1 | 0 | 0.30 |
| Zimbabwe | 8 | 2 | 1 | 3.50 | 1 | 0 | 0.21 |
| Colombia | 32 | 1 | 0 | 3.47 | 0 | 0 | 1.41 |
| Mozambique | 7 | 2 | 0 | 3.45 | 0 | 0 | 0.23 |
| India | 13 | 2 | 0 | 3.45 | 2 | 0 | 0.51 |
| Mexico | 23 | 1 | 0 | 3.39 | 0 | 0 | 0.54 |
| Cuba | 22 | 1 | 0 | 3.26 | 0 | 0 | 0.36 |
| Indonesia | 15 | 1 | 0 | 3.14 | 1 | 1 | 0.40 |
| Belize | 16 | 1 | 0 | 3.03 | 0 | 0 | 0.42 |
| eSwatini | 6 | 2 | 0 | 2.94 | 0 | 0 | 0.07 |
| Guatemala | 13 | 1 | 0 | 2.83 | 0 | 0 | 0.44 |
| Nicaragua | 15 | 1 | 0 | 2.82 | 0 | 0 | 0.45 |
| Honduras | 16 | 1 | 0 | 2.80 | 0 | 0 | 0.46 |
| Costa Rica | 17 | 1 | 0 | 2.70 | 0 | 0 | 0.98 |
| Trinidad and Tobago | 13 | 1 | 0 | 2.62 | 1 | 1 | 0.74 |
| Liberia | 10 | 2 | 1 | 2.58 | 0 | 0 | 0.18 |
| Burkina Faso | 11 | 1 | 0 | 2.51 | 0 | 0 | 0.53 |
| Benin | 18 | 1 | 0 | 2.50 | 0 | 0 | 0.61 |
| Peru | 24 | 1 | 0 | 2.45 | 1 | 0 | 1.53 |
| Eq. Guinea | 9 | 2 | 1 | 2.42 | 1 | 0 | 0.31 |
| Uganda | 11 | 1 | 0 | 2.36 | 0 | 0 | 0.27 |
| Botswana | 8 | 1 | 0 | 2.27 | 0 | 0 | 0.08 |
| Togo | 12 | 1 | 0 | 2.16 | 0 | 0 | 0.56 |
| Dominican Rep. | 8 | 2 | 0 | 2.08 | 1 | 0 | 1.01 |
| United States of America | 14 | 1 | 0 | 2.06 | 0 | 0 | 0.12 |
| Malawi | 6 | 1 | 1 | 2.01 | 0 | 0 | 0.21 |
| Taiwan | 8 | 1 | 0 | 1.95 | 0 | 0 | 0.09 |
| Sri Lanka | 6 | 1 | 0 | 1.84 | 0 | 0 | 0.04 |
| Argentina | 12 | 1 | 0 | 1.83 | 0 | 0 | 0.39 |
| Puerto Rico | 11 | 1 | 0 | 1.77 | 0 | 0 | 0.25 |
| Sierra Leone | 8 | 1 | 1 | 1.68 | 0 | 0 | 0.10 |
| Panama | 15 | 0 | 0 | 1.65 | 1 | 0 | 0.96 |
| Congo | 11 | 1 | 0 | 1.64 | 1 | 0 | 0.33 |
| Jamaica | 6 | 1 | 0 | 1.55 | 0 | 0 | 0.16 |
| Kenya | 5 | 1 | 0 | 1.48 | 0 | 0 | 0.17 |
| Paraguay | 12 | 0 | 0 | 1.47 | 0 | 0 | 0.40 |
| Central African Rep. | 10 | 0 | 0 | 1.31 | 0 | 0 | 0.16 |
| Fiji | 2 | 1 | 0 | 1.29 | 0 | 0 | 0.01 |
| Tonga | 2 | 1 | 0 | 1.29 | 0 | 0 | 0.01 |
| Suriname | 17 | 0 | 0 | 1.25 | 0 | 0 | 0.70 |
| Rwanda | 2 | 1 | 0 | 1.24 | 0 | 0 | 0.02 |
| Bahamas | 1 | 1 | 0 | 1.22 | 0 | 0 | 0.01 |
| Timor-Leste | 1 | 1 | 0 | 1.22 | 0 | 0 | 0.01 |
| Cayman Is. | 1 | 1 | 0 | 1.22 | 0 | 0 | 0.01 |
| Ecuador | 16 | 0 | 0 | 1.17 | 0 | 0 | 0.47 |
| Guinea-Bissau | 5 | 0 | 0 | 1.00 | 0 | 0 | 0.09 |
| Niger | 2 | 0 | 0 | 0.76 | 0 | 0 | 0.03 |
| S. Sudan | 1 | 1 | 0 | 0.74 | 1 | 0 | 0.10 |
| New Caledonia | 4 | 1 | 0 | 0.72 | 2 | 1 | 0.28 |
| Chad | 4 | 0 | 0 | 0.50 | 0 | 0 | 0.15 |
| Japan | 6 | 0 | 0 | 0.49 | 0 | 0 | 0.07 |
| Mali | 7 | 0 | 0 | 0.48 | 0 | 0 | 0.47 |
| Lesotho | 3 | 0 | 0 | 0.43 | 0 | 0 | 0.01 |
| Senegal | 9 | 0 | 0 | 0.41 | 0 | 0 | 0.50 |
| Nepal | 3 | 0 | 0 | 0.41 | 0 | 0 | 0.03 |
| Burundi | 5 | 0 | 0 | 0.40 | 0 | 0 | 0.16 |
| Myanmar | 2 | 0 | 0 | 0.40 | 0 | 0 | 0.02 |
| El Salvador | 5 | 0 | 0 | 0.34 | 0 | 0 | 0.06 |
| Vietnam | 5 | 0 | 0 | 0.32 | 0 | 0 | 0.07 |
| Papua New Guinea | 3 | 0 | 0 | 0.25 | 0 | 0 | 0.02 |
| Cambodia | 3 | 0 | 0 | 0.23 | 1 | 0 | 0.08 |
| Seychelles | 2 | 0 | 0 | 0.19 | 0 | 0 | 0.04 |
| Mauritius | 1 | 0 | 0 | 0.19 | 1 | 0 | 0.06 |
| Vanuatu | 2 | 0 | 0 | 0.15 | 0 | 0 | 0.01 |
| Laos | 2 | 0 | 0 | 0.09 | 0 | 0 | 0.02 |
| Uruguay | 1 | 0 | 0 | 0.08 | 0 | 0 | 0.00 |
| Malaysia | 1 | 0 | 0 | 0.02 | 0 | 0 | 0.01 |
| French Guiana | 0 | 0 | 0 | 0.00 | 0 | 0 | 0.00 |

**Appendix 5.** Top 50 ecoregions in which *Scleria* is present ranked by their sum of EDGE2 scores. Richness: number of species present, N: number of EDGE2 and EcoDGE species, Main: number of species included in the main EDGE2 and EcoDGE lists, Sum: sum of EDGE2 and EcoDGE scores.

| **Ecoregion** | **Richness** | **EDGE2** | | | **EcoDGE** | | |
| --- | --- | --- | --- | --- | --- | --- | --- |
|  |  | **N** | **Main** | **Sum** | **N** | **Main** | **Sum** |
| Central Zambezian Miombo woodlands | 39 | 8 | 7 | 15.32 | 3 | 0 | 2.04 |
| Madagascar subhumid forests | 22 | 7 | 3 | 9.50 | 2 | 0 | 1.09 |
| Cerrado | 45 | 3 | 1 | 7.09 | 2 | 0 | 1.72 |
| East Sudanian savanna | 22 | 5 | 1 | 5.76 | 1 | 0 | 0.85 |
| Llanos | 24 | 3 | 1 | 5.68 | 0 | 0 | 0.86 |
| Guinean forest-savanna mosaic | 28 | 4 | 1 | 5.61 | 1 | 0 | 0.81 |
| Madagascar lowland forests | 16 | 2 | 1 | 5.48 | 1 | 0 | 0.65 |
| Northwestern Congolian lowland forests | 20 | 4 | 1 | 5.28 | 2 | 0 | 0.37 |
| Zambezian and Mopane woodlands | 15 | 3 | 1 | 5.28 | 0 | 0 | 0.18 |
| Mato Grosso seasonal forests | 28 | 2 | 1 | 4.87 | 1 | 0 | 1.05 |
| Western Congolian forest-savanna mosaic | 18 | 3 | 1 | 4.87 | 3 | 0 | 0.80 |
| Tenasserim-South Thailand semi-evergreen rain forests | 13 | 3 | 1 | 4.82 | 0 | 0 | 0.19 |
| Zambezian flooded grasslands | 13 | 2 | 0 | 4.76 | 0 | 0 | 0.24 |
| Eastern Arc forests | 14 | 3 | 1 | 4.70 | 1 | 0 | 0.39 |
| South China-Vietnam subtropical evergreen forests | 12 | 3 | 0 | 4.70 | 0 | 0 | 0.19 |
| Guianan savanna | 21 | 2 | 0 | 4.61 | 0 | 0 | 0.73 |
| Drakensberg montane grasslands, woodlands and forests | 13 | 2 | 0 | 4.33 | 0 | 0 | 0.11 |
| Eastern Guinean forests | 21 | 2 | 0 | 4.28 | 0 | 0 | 0.70 |
| West Sudanian savanna | 21 | 2 | 0 | 4.23 | 0 | 0 | 0.64 |
| Western Guinean lowland forests | 20 | 3 | 2 | 4.15 | 0 | 0 | 0.69 |
| Southern Rift montane forest-grassland mosaic | 14 | 2 | 1 | 3.94 | 0 | 0 | 0.29 |
| Angolan Miombo woodlands | 11 | 2 | 1 | 3.90 | 1 | 0 | 0.20 |
| Cardamom Mountains rain forests | 11 | 3 | 1 | 3.89 | 1 | 0 | 0.18 |
| Southern Congolian forest-savanna mosaic | 15 | 2 | 1 | 3.83 | 1 | 0 | 0.52 |
| Guinean montane forests | 17 | 3 | 2 | 3.52 | 0 | 0 | 0.22 |
| Petén-Veracruz moist forests | 21 | 1 | 0 | 3.52 | 0 | 0 | 0.54 |
| Eastern Miombo woodlands | 9 | 2 | 0 | 3.29 | 0 | 0 | 0.22 |
| Alto Paraná Atlantic forests | 23 | 1 | 0 | 3.28 | 0 | 0 | 0.77 |
| Central Indochina dry forests | 11 | 2 | 0 | 3.22 | 0 | 0 | 0.15 |
| Bahia coastal forests | 27 | 1 | 0 | 3.21 | 2 | 2 | 1.82 |
| Atlantic Equatorial coastal forests | 16 | 2 | 1 | 3.20 | 1 | 0 | 0.39 |
| Arnhem Land tropical savanna | 18 | 1 | 0 | 3.20 | 0 | 0 | 0.18 |
| Southern Miombo woodlands | 9 | 2 | 2 | 3.19 | 1 | 0 | 0.35 |
| Southern Africa bushveld | 6 | 2 | 0 | 3.05 | 0 | 0 | 0.07 |
| Guianan piedmont and lowland moist forests | 22 | 1 | 0 | 3.03 | 0 | 0 | 0.85 |
| Queensland tropical rain forests | 15 | 1 | 0 | 2.99 | 1 | 0 | 0.24 |
| Cape York Peninsula tropical savanna | 16 | 1 | 0 | 2.96 | 0 | 0 | 0.17 |
| Central American pine-oak forests | 18 | 1 | 0 | 2.94 | 0 | 0 | 0.47 |
| Western Java rain forests | 12 | 1 | 0 | 2.93 | 1 | 1 | 0.36 |
| KwaZulu-Cape coastal forest mosaic | 12 | 1 | 0 | 2.91 | 0 | 0 | 0.10 |
| Caqueta moist forests | 21 | 1 | 0 | 2.90 | 0 | 0 | 0.72 |
| Maputaland coastal forest mosaic | 10 | 1 | 0 | 2.90 | 0 | 0 | 0.11 |
| Pantanal | 14 | 1 | 0 | 2.81 | 0 | 0 | 0.39 |
| Southeastern Indochina dry evergreen forests | 7 | 2 | 1 | 2.80 | 0 | 0 | 0.09 |
| Bolivian Yungas | 18 | 1 | 0 | 2.69 | 0 | 0 | 0.57 |
| Kimberly tropical savanna | 13 | 1 | 0 | 2.68 | 0 | 0 | 0.13 |
| Einasleigh upland savanna | 12 | 1 | 0 | 2.65 | 0 | 0 | 0.12 |
| Tocantins/Pindare moist forests | 18 | 1 | 0 | 2.59 | 0 | 0 | 0.55 |
| Southern Pacific dry forests | 12 | 1 | 0 | 2.58 | 0 | 0 | 0.23 |
| Chiquitano dry forests | 26 | 0 | 0 | 2.57 | 0 | 0 | 0.83 |
